# An investigation of the relationship between changes in psychosocial domains with a virtual salsa class and personality, perceived performance and enjoyment in neurotypical adults

**DOI:** 10.64898/2026.02.22.707137

**Authors:** Rimsha Amin, Sergiu-Gabriel Duplea, Marina Gadalla, Jeremy Pullara, Andrew Lam, Callum Smith, Hayley Ng, Kara K Patterson

## Abstract

This pre-post-test study investigated 1) pre-post changes in psychosocial domains with a single virtual salsa class; 2) effect sizes relative to an in-person class, and 3) individual factors, including personality, perceived performance, and enjoyment. An experimental group (n=33) of novice dancers 18-30 years old, participated in a single virtual salsa class. Positive and Negative Affect Scale (PAS, NAS), Perceived Stress Scale (PSS), and the Inclusion in Community and Self-Scale (ICS) were administered before and after class. Participants completed the Big Five Inventory-10 (BFI-10) before, and rated their performance and enjoyment (ordinal scale 1-5) after class. Effect sizes were calculated, and pre-post changes were analyzed with Wilcoxon signed-rank tests. Relationships between pre-post changes and individual factors were analyzed with Spearman’s rank correlations. PAS, NAS, PSS, and ICS significantly improved and effect sizes were larger than those for an in-person salsa class except for ICS. Change in NAS was negatively correlated with neuroticism. These results suggest that a virtual salsa class may improve mood, stress, and social connection similar to in-person classes and change in mood may be influenced by personality traits such as neuroticism. Understanding the psychosocial effects of virtual dance and the influence of individual factors will facilitate implementation of dance as an accessible rehabilitation intervention to improve psychosocial well-being.

## INTRODUCTION

Dance is gaining recognition as a complementary intervention to conventional rehabilitation. Clinicians appreciate its potential as a holistic intervention that improves both physical and psychosocial well-being.(Wilberforce, 2025). Physical benefits include improved cardiorespiratory fitness, mobility, and balance.(Hui et al., 2009) Psychosocial benefits include reduced perceived stress, increased social connection, and improved mood.(Quiroga et al., 2010; West et al., 2004) These benefits appear consistent across various dance styles, and people who dance experience these benefits regardless of skill level, age, or clinical condition.(Fong Yan et al., 2018) However, further investigation is needed to advance the use of dance in rehabilitation, which can be considered a complex intervention as per the Medical Research Council. (Council, 2019) A randomized controlled trial is required to establish the effectiveness of an intervention. But, in the case of complex interventions, the MRC framework asserts that another critical step is understanding the “active ingredients” or the physiological mechanisms through which the intervention has its impact (Council, 2019). Understanding the active ingredients of a complex intervention can inform the design of an RCT and facilitate the selection of an appropriate control intervention. The mechanisms underlying the physical and psychosocial benefits of dance are largely unexplored.

In our previous work, we suggested two (of likely many) potential active ingredients of dance interventions: the synchronization of movement with music and with other dancers.(Scott et al., 2025) Pre-post changes in mood, stress, and social connection are possible with a single dance class.(Scott et al., 2025) Furthermore, preliminary evidence suggests that changes in mood are associated with how well a person synchronizes their movements with their dance partner.(Scott et al., 2025) It is important to note that this previous work focused on in-person dance classes. It is unclear whether virtual dance classes provide similar psychosocial benefits, given that the format may not provide the same opportunity to synchronize movement.^15,16^ Furthermore, others have identified unique aspects of in-person dance practice, including physical proximity and tactile cues, which are missing in virtual classes. (Rugh et al., 2024) COVID necessitated moving services and activities (including rehabilitation and dance classes) online. Although in-person delivery has resumed, online delivery remains a viable option. Therefore, it is important to determine whether virtual dance classes are associated with pre-post changes in mood, stress, and connection that are comparable to those in in-person classes.

In addition to identifying active ingredients of a complex intervention, it is equally important to characterize individual factors that may influence the outcomes a person experiences with that intervention. Potential individual factors that could influence dance outcomes include personality traits, perceived performance in the class, and enjoyment of the class. Different personality dimensions are associated with specific movement characteristics when dancing to music.(Luck et al., 2010) For example, extroversion is associated with faster movement speed of the head, hands, and center of mass.(Luck et al., 2010) Neuroticism is associated with jerky movements (rate of change of acceleration) of the same body parts.(Luck et al., 2010) Thus, personality traits may influence the outcomes people achieve with dance by affecting how well they synchronize their movement with music and other members of the class. Finally, perceived performance and enjoyment of the class could also influence dance outcomes. The mastery hypothesis suggests that successfully completing an effortful task fosters a sense of accomplishment and enhances mood(Bartholomew & Miller, 2002). An investigation of an aerobic dance class found that changes in positive affect were moderated by ratings of perceived performance and enjoyment of the class.^18^ Thus, it is possible that how a person perceives their performance in the dance class, and their enjoyment of the class may influence the impact the dance class has on their psychosocial well-being.

A deeper understanding of the psychosocial effects of a virtual dance class and the influence of individual factors on those effects will facilitate future work on implementing dance as a complex rehabilitation intervention. For example, if we learn that certain personality traits influence the effects of dance on stress or mood, then screening people for those traits would inform clinical decision-making about whether dance is an appropriate intervention.

In summary, virtual dance classes may serve as an accessible, holistic rehabilitation intervention. To implement such an intervention, an important first step is to determine whether the immediate effects on mood, stress, and connection are similar to those observed with in-person classes. Equally important to understand are the individual factors that may influence those dance outcomes. Thus, the primary objective of this study is to investigate the pre- post- changes in mood, perceived stress, and social connection associated with a single virtual beginner salsa class for neurotypical adults and to compare the effect sizes to a similar in-person dance class. A second objective is to examine the relationship between changes in psychosocial domains from the salsa class and individual factors, including personality, perceived performance, and perceived enjoyment.

## MATERIALS AND METMETHODS

This was a cross-sectional study with a pre- and post-test design, approved by the University of Toronto’s Health Sciences Research Ethics Board (protocol #43663).

### Participants

Neurotypical adults between the ages of 18 – 30 years were recruited from the community using posters and social media advertising. Individuals were eligible to participate if they had internet access and a webcam and were able to follow multi-step instructions in English. Exclusion criteria included the presence of neurological or musculoskeletal conditions, impairments in balance, mobility, or hearing that could compromise safety or independent participation, and >1 year of formal salsa dance experience. Since one of the variables of interest was social connection, an additional eligibility criterion was that participants did not have more than one close relationship (beyond a passing acquaintance) with anyone in the dance class they attended. All participants provided informed consent via an electronic consent form.

### Dance Classes

Each participant attended a single, 45-minute, beginner salsa lesson delivered via Zoom. Participants were assigned to one of six dance classes based on the availability they provided to the research team and were provided with the names of the other participants in their class. If a participant identified more than one individual with whom they had a close relationship, they were reassigned to another dance class. Before the class, participants received the following instructions via email: 1) ensure they had a safe, obstacle-free, open space that was approximately twice their arm span, 2) wear proper footwear, 3) dress comfortably, 4) keep their camera on throughout the duration of the class, and 5) position the camera such that their full body was visible to facilitate instructor feedback. All six classes were taught by a professional dance instructor with 15 years of experience teaching both in-person and virtual salsa classes. The music, instructions, and content were the same across the six classes. Each class began with instruction of the basic salsa steps and progressed to multi-step salsa dance moves by the end of the class. The instructor provided verbal feedback and visual demonstrations to participants as needed.

### Data Collection Protocol

All variables of interest were collected using electronic questionnaires (Qualtrics, Seattle, USA). Links to two of the questionnaires were emailed to participants to be completed prior to the dance class: a sociodemographic questionnaire and the abbreviated Big Five Inventory-10 (BFI-10) (Rammstedt & John, 2007) to assess personality. Participants were asked to join their assigned dance class 10 –15 minutes prior to the start time and remain in the Zoom room 10 – 15 minutes following the class to complete the pre- and post-test questionnaires to measure dance outcomes (mood, stress, social connection) and individual factors (enjoyment, performance). The link to these test questionnaires was provided to the participants via the Zoom chat.

### Variables of Interest

Mood was measured using the Positive and Negative Affect Schedule (PANAS). This scale consists of 20 items, with 10 items measuring positive affect (PAS) and 10 items measuring negative affect (NAS).(Watson et al., 1988) The PAS and NAS items are scored separately, with higher scores indicating a more positive or more negative effect, respectively.(Watson et al., 1988) Perceived stress was measured using the Perceived Stress Scale (PSS).(Cohen et al., 1983) The PSS is a 10-item scale that is designed to measure how stressful situations in one’s life are perceived to be, with higher scores indicating greater levels of perceived stress.(Cohen et al., 1983) Social connection to the dance class was measured using the Inclusion of Community in Self Scale (ICS) which is a meaningful measure of community connectedness distinct from close relationship connectedness.(Mashek et al., 2007) The ICS is a one-item ordinal scale comprised of a series of six images of two increasingly overlapping circles, with one circle representing the self, and the other representing the community. Each image was assigned a numerical value corresponding to the level of social connectedness for subsequent statistical analysis, with 1 = least perceived social connectedness (circles completely separate) and 6= the most perceived social connectedness (circles completely overlapped).(Mashek et al., 2007) Participants were instructed to select the image that best illustrates their relationship to others in the dance class.

Individual factors of interest included personality, perceived performance, and perceived enjoyment. Personality with the Big Five Inventory-10 (BFI-10). This measure consists of 10 items that assess five personality traits: extraversion, agreeableness, neuroticism, conscientiousness, and openness to experience.(Rammstedt & John, 2007) Perceived performance and perceived enjoyment were assessed following the salsa class using five-point ordinal scales; 1 = “very little” or “very poor,” to 5= “very much” or “very well.”(Bartholomew & Miller, 2002)

### Data Analysis

Questionnaire data were exported from Qualtrics to Microsoft Excel for further processing and analysis. PAS, NAS, PSS, ICS and BFI-10 scores for neuroticism and extraversion were calculated in Excel according to the guidelines for each measure. All statistical analyses were performed in SAS 9.4 (SAS Institute Inc., USA). Demographics, individual factors, and pre- and post-class primary outcome values were summarized with descriptive statistics for the whole study group and each salsa class. Continuous data were summarized with mean and standard deviation (sd), and ordinal data were summarized with median and 1^st^ and 3^rd^ quartiles [Q1, Q3]. The six salsa classes were examined for similarity at baseline in demographics, PAS, NAS, PSS, ICS, BFI scores for extraversion and neuroticism, and ratings of perceived performance and enjoyment using one-way ANOVAs (ordinal data were rank-transformed prior to the ANOVA). Fisher’s exact tests were used to compare the six classes on proportions of participants who had previous dance training and proportions of participants who identified as men and women. The primary study objective (examine pre-post changes in mood, stress and social connection and compare to an in-person class) was addressed using Wilcoxon signed-rank tests on change scores for PAS, NAS, PSS and ICS. Effect sizes were also calculated as r = Z/√n (Tomczak & Tomczak, 2014) and compared to effect sizes previously published for an in-person salsa class.(Scott et al., 2025) It is important to note that the in-person salsa class was taught by the same dance instructor in the present study, and the outcome measures were identical to the present study.(Scott et al., 2025). The second objective (characterize relationships between changes in primary outcomes and individual factors) was addressed with Spearman’s rank correlations to quantify the association between change scores for the dance outcomes (PAS, NAS, PSS, ICS) and the individual factors (perceived performance, enjoyment and BFI scores for extroversion, neuroticism, agreeableness, conscientiousness, and openness). Significance level was set to P<.05.

## RESULTS

Forty-nine neurotypical young adults (women n=46, men n=3) contacted the research group to participate in the study. Sixteen participants (women n=15, men n=1) did not respond after the initial email from the research group. Thirty-three people met the eligibility criteria (women n=31; men n=2) and completed the dance class and study outcome measures. The demographics and ratings for performance and enjoyment by the whole group and each salsa class are summarized in Table 1. There were no significant differences among the 6 classes for demographics, baseline values of the primary outcomes, median ratings of perceived performance and enjoyment, or BFI scores for extraversion and neuroticism.

**Table 1.**
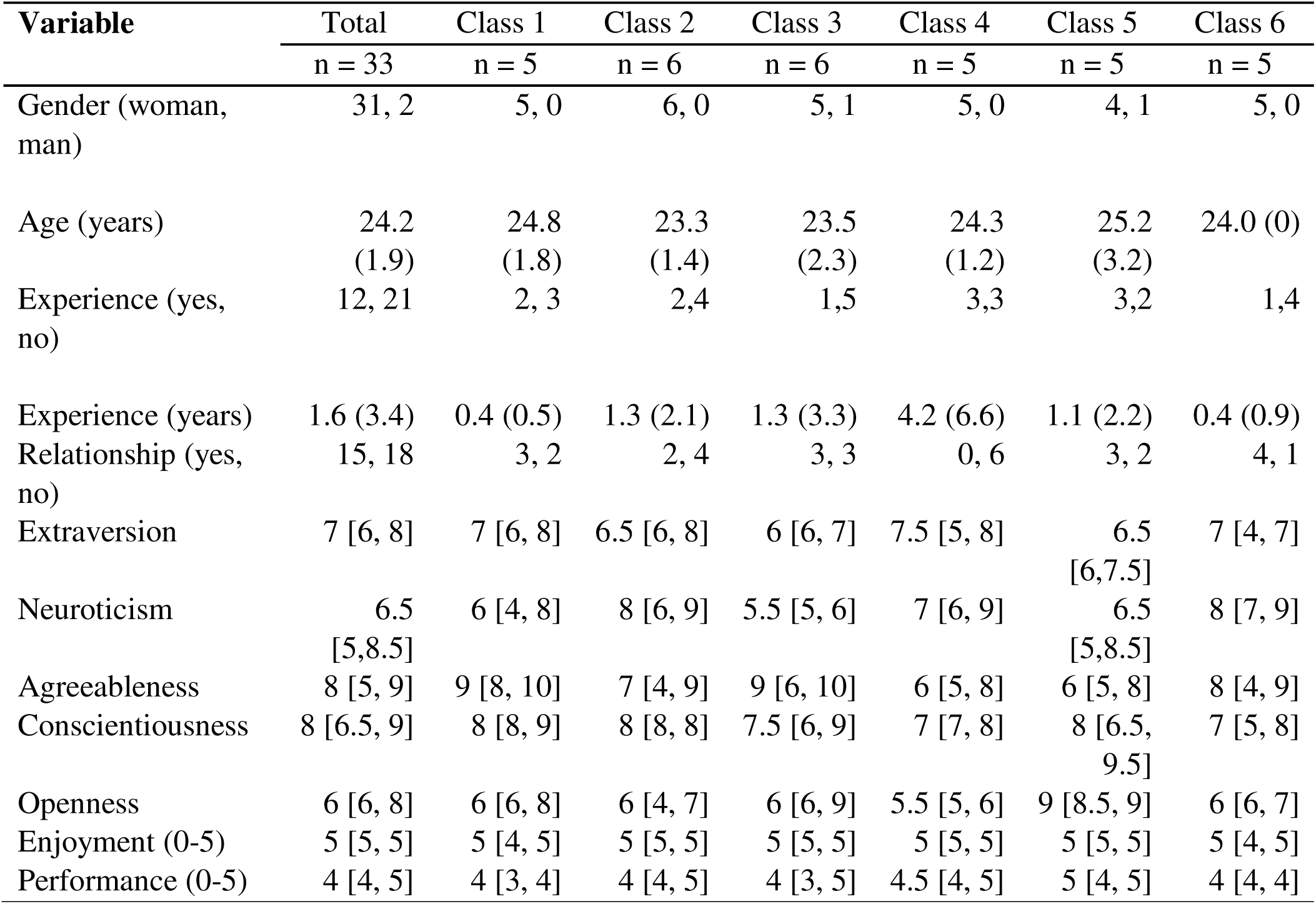
Demographics for the whole group and each salsa class. Values are presented as mean (sd) for continuous variables and median [Q1, Q3] for ordinal variables.

The median [Q1, Q3] pre- and post-values for PAS, NAS, PSS, and ICS and effect sizes are summarized for the whole group in Table 2.

**Table 2.** Outcome measures pre- and post-virtual salsa class. Values are presented as median [Q1, Q3].

| Outcome Measure | Pre class | Post class | Effect size | In-person class effect size(Scott et al., 2025) |
| --- | --- | --- | --- | --- |
| PAS* | 28 [23, 34] | 42 [34, 43] | -0.86 | -0.75 |
| NAS* | 13 [12, 21] | 11 [10, 12] | -0.74 | -0.52 |
| PSS* | 20 [17, 25] | 18 [15, 22] | -0.75 | -0.56 |
| ICS* | 2 [1, 3] | 3 [2, 4] | -0.70 | -0.82 |

Abbreviations: PAS = Positive Affect Scale; NAS = Negative Affect Scale; PSS: Perceived Stress Scale; ICS = The Inclusion of Community in Self Scale. \* p<0.0001

All variables were significantly different pre- to post-salsa class. Spearman’s rank correlation coefficients for the relationships between change in outcomes of interest (PAS, NAS, PSS, ICS) and individual factors (extraversion, neuroticism, perceived performance, and perceived enjoyment) are presented in Table 3.

**Table 3.** Correlation coefficients for relationships between individual factors and change scores for outcome measures.

| Variable | Performance | Enjoyment | Extraversion | Neuroticism |
| --- | --- | --- | --- | --- |
| PAS change score | -0.24 | -0.02 | -0.30 | 0.35 |
| NAS change score | 0.18 | 0.10 | 0.08 | -0.46* |
| PSS change score | 0.20 | -0.06 | -0.04 | -0.07 |
| ICS change score | 0.06 | 0.01 | 0.07 | 0.19 |

Abbreviations: PAS = Positive Affect Scale; NAS = Negative Affect Scale; PSS = Perceived Stress Scale; ICS = The Inclusion of Community in Self Scale, \* $p=0.0088$

Figure 1 illustrates a statistically significant (p= 0.0088) negative correlation between neuroticism and change in NAS. All other relationships did not show a statistically significant correlation.

**Figure 1.**
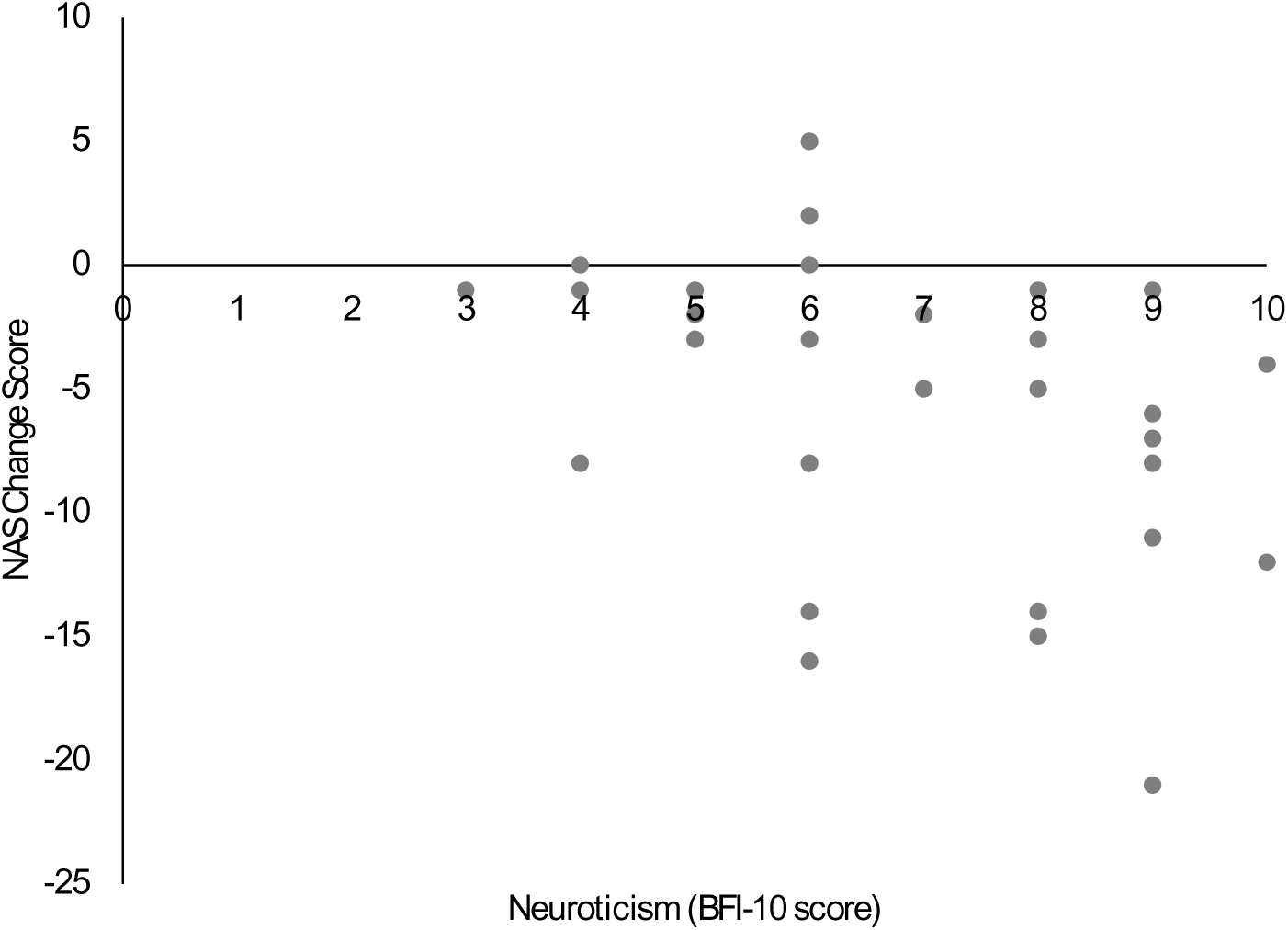
Relationship between neuroticism and change in negative affect after a single virtual salsa dance class. Negative change scores in Negative Affect Scale (NAS) represent an improvement. Markers represent individual participants (n=32, missing data for BFI-10 n=1)

## DISCUSSION

The main findings of this study are that a virtual salsa class was associated with pre-post changes in mood, perceived stress, and social connection, and the magnitude of change in negative affect was associated with the personality trait of neuroticism. The more neurotic a participant was, the greater the change in negative affect they experienced with the salsa class. These results demonstrate that psychosocial benefits are possible with virtual dance classes. The results also lend support to our proposed model of the active ingredients of dance and potential influencing factors.

The present study compared effect sizes for the virtual beginner salsa class with those from a previous study we conducted on an in-person beginner salsa class, using the exact same outcome measures (i.e., PANAS, PSS, ICS).(Scott et al., 2025) The same salsa instructor conducted all classes in both studies, thereby ensuring consistency in certain variables between the two studies, such as teaching style and instructor personality. This allows for some conclusions to be drawn about the impact of virtual delivery of the salsa class. First, the effect sizes for all outcome measures were large for both the present virtual and previous in-person classes. Interestingly, compared to the in-person class, the virtual salsa class had larger effect sizes for PAS, NAS, and PSS. Second, as proposed, the virtual class produced a smaller effect size for social connection as measured by ICS. This may be explained by the absence of physical proximity, physical contact with dance partners, and synchronization of movement with other dancers during partner work in the virtual class format. Third, these results lend indirect support to the proposed model for active ingredients of dance. Specifically, that synchronized movement is one potential underlying mechanism of the effects of dance. Future work should explore this mechanism with direct measures of dance movements. Fourth, it is worth noting that the large effect sizes for both classes indicate that either virtual or in-person delivery are effective at improving mood, stress, and social connection. Other factors, such as personality, could then determine which delivery mode is most appropriate for an individual.

A unique finding of the present study is the relationship between personality and the benefits derived from the virtual salsa class. Specifically, participants with a greater degree of neuroticism, as measured by the BFI-10, experienced larger decreases in negative affect with the salsa class. These results inform another component of our proposed framework for dance as a complex rehabilitation intervention. Specifically, personality traits may interact with how dance exerts its effect on mood. Personality has been associated with different characteristics of music-induced movement (Luck et al., 2010). Neuroticism is associated with “jerky and accelerated movement of the head, hands, feet and centre of mass”(Luck et al., 2010). Our framework posits that dance-movement characteristics associated with certain personality traits may influence synchronization of movement with music and with other dancers in the class, which in turn could affect the psychosocial benefits derived from the class. Alternatively, emotional reactivity to negative-mood induction may have contributed to the observed relationship between neuroticism and change in NAS. People with higher levels of neuroticism experience intense negative emotions(Kalokerinos et al., 2020) and they are more responsive to situations designed to induce a negative mood.(Larsen & Ketelaar, 1991) In the present work, the reverse seemed to occur, one of the goals of the dance class was to reduce negative mood and people with higher scores in neuroticism exhibited the largest improvement in negative affect.

Other personal factors, such as enjoyment of the dance class and perceived performance, were not associated with changes in mood, stress, or social connection. This contrasts with previous work investigating changes in mood with a virtual dance class featuring jazz, ballet, and contemporary styles.(Rugh et al., 2024) In this previous work, self-reports of enjoyment, rated on a scale from 0 to 10, was significantly correlated with change in positive affect measured by PANAS (r = .351).(Rugh et al., 2024) In the present study a smaller scale was used to rate enjoyment (i.e., 0-5) and most participants rated enjoyment at the top of the scale (i.e. 4 or 5) whereas in the previous study responses ranged from 5 to 10. The limited range of enjoyment (and performance ratings) in the present study may have hampered the ability to detect relationships with changes in mood, stress, and social connection.

It is important to note that 94% of the study group identified as women. Previous work with a large sample size (n= 1,003) suggests that the influence of demographics on the PANAS, such as age, education, and gender can be disregarded. (Crawford & Henry, 2004) However, women consistently report higher levels of perceived stress compared to men.(Kneavel, 2021) Furthermore, a meta-analysis revealed that affect induction methods have stronger effect sizes for women compared to men,(Joseph et al., 2020) suggesting that in the present study, the effect size for dance on positive and negative affect may have been different if the sample had included more men. Additionally, the previous work with the in-person salsa class had more men (38%), which may have contributed to the observed differences in effect sizes for positive and negative affect from the current effect sizes for the virtual salsa class.

This study had some limitations. First, as mentioned above, most of the study participants were female, which may limit the generalizability of our findings. Subsequent studies should strive for a sample that is more gender diverse, which would facilitate sex and gender-based analysis of the psychosocial benefits of virtual dance classes. Second, as with all studies that use self-report questionnaires, recall bias may have influenced our results. In particular, the relatively short interval between completing the pre- and post- surveys (∼3 hours) means participants could have remembered their answers to the pre-class questionnaires when completing their post-class questionnaires. However, previous work on dance(Rugh et al., 2024) used a similar time interval, as it was noted that the acute physiological effects of exercise are most prominent up to 120 minutes after completion of the activity. Furthermore, the participants were blinded to the hypotheses of this study which may have mitigated the ‘good participant’ effect.(Nichols & Maner, 2008) Third, the initial exploratory nature of this pre- and post-design study means there was no control group. Thus, definitive conclusions about the psychosocial benefits of salsa dance cannot be made.

## CONCLUSION

Dance is gaining recognition as a holistic rehabilitation intervention, demonstrating both physical and psychosocial benefits. During the COVID-19 pandemic, exercise and dance classes for rehabilitation purposes were forced to pivot to an online format. While the capacity for physical improvements with virtual classes seemed apparent, the capacity to improve psychosocial well-being was uncertain. The present study suggests that virtual dance has large effect sizes for mood, stress, and social connection despite the lack of in-person contact. The inverse relationship between changes in negative mood and neuroticism highlights the need to account for personality in future investigations on the clinical application of dance. Future work should include a randomized controlled study to investigate changes in psychosocial well-being with repeated virtual salsa classes, while accounting for the potential influence of individual factors such as neuroticism in a gender diverse group.

## Acknowledgements

This research was completed in partial fulfillment of the requirements for a MScPT degree at the University of Toronto. The authors of this study would like to acknowledge the contributions of the participants of this research study.

## Data availability

The datasets presented in this article are not readily available because REB approval was not obtained for this type of data usage and/or dissemination and the participants of this study did not give written consent for their individual data to be shared publicly.

## Author contributions (CrediT taxonomy)

**Rimsha Amin, Sergiu-Gabriel Duplea, Marina Gadalla, Jeremy Pullara, Andrew Lam, and Callum Smith** were involved in data curation, formal analysis, investigation, methodology, visualization, writing-original draft, writing-review & editing.

**Hayley Ng** was involved in data curation, investigation, methodology, supervision and writing-review & editing.

**Kara K Patterson** was involved in conceptualization, formal analysis, funding acquisition, methodology, project administration, resources, supervision, visualization, and writing-review & editing.

All authors have reviewed and approved the version of the manuscript submitted and agree to be accountable for all aspects of the work.

